# Wavelet Transform improves time-frequency characterization of the Electroretinographic Oscillatory Potentials as compared to Short-Time Fourier Transform across experimental and clinical models

**DOI:** 10.64898/2026.09.26.754708

**Authors:** Patrick Reiman, Sean Ghiam, Avigail Beryozkin, Valentina Stein Uman, Ayala Sahel, Jean-Francois Mouillet, Lucas Janischetz-Kriegl, Jordan Safran, Eitan Shemuelian, Oliver Beale, Samer Khateb, Anfisa Ayalon, Richard Hagan, Alkiviades Liasis, Dimitrios Stavropoulos, Kun-Che Chang, Takaki Kuwajima, Kamakshi Mehta, Issam Aldiri, Avi Caspi, Isabelle Audo, Eyal Banin, Jose-Alain Sahel, Boris Rosin

## Abstract

**Purpose:** To compare continuous wavelet transform (CWT) and short-time Fourier transform (STFT) analyses of the oscillatory potentials (OPs) of the dark-adapted full-field electroretinogram (FFERG).

**Methods:** FFERG recordings from Royal College of Surgeons (RCS) and healthy Brown Norway (BN) rats, wild-type and Atoh7 enhancer-knockout (KO) mice, and patients with diabetic retinopathy and healthy controls were analyzed with STFT and with CWT employing the real Morlet wavelet, after 60 Hz high-pass and sharp 60 Hz notch filtering. Time- and frequency-localized OP features were quantified and the methods compared on detection of transient OP signals. Both were additionally validated using synthetic OP-like bursts of known frequency embedded in each cohort’s recorded noise.

**Results:** CWT revealed distinct time–frequency patterns and disease-specific OP alterations across all rat, mouse, and human cohorts, with improved temporal localization and frequency resolution, whereas STFT merged transient features and was more susceptible to noise. CWT-derived dominant-frequency differences reached significance in all three models; the corresponding STFT difference reached significance only in rats (separate tests; methods not directly compared). In synthetic bursts, both methods recovered injected frequencies to within 1–2.5 Hz, but only CWT resolved two bursts separated by 15 ms.

**Conclusions:** CWT employing the real Morlet wavelet provides a robust, physiologically relevant evaluation of OPs, with improved temporal localization and reduced spectral dispersion relative to STFT in clinical and experimental settings.

**Translational Relevance:** CWT analysis may improve the characterization of inner retinal dysfunction by preserving transient OP time–frequency structure, as illustrated in preclinical rodent models and diabetic retinopathy patients.

## Introduction

Electroretinography (ERG) is a widely used method for evaluating retinal function. The full-field electroretinogram (FFERG) is a noninvasive electrophysiological test quantifying retinal electrical activity in response to controlled diffuse flashes of light (1). The resultant response is a complex waveform which consists of several superimposed waves, reflecting the electrical activity of different retinal cells and layers (2). One of these components is the oscillatory potentials (OPs), which appear on the ascending edge of the b-wave in scotopic and photopic bright-flash recordings (3, 4).

Although the exact cellular origin of OPs is not firmly established, studies suggest they reflect complex inner retinal interactions, such as membrane voltage oscillations along the bipolar cell axis induced by inhibitory feedback from amacrine cells (3, 5). As such, OPs offer unique insights into inner retinal function and a means for its objective quantification. They have demonstrated clinical utility in various retinal diseases involving the inner retina, including retinitis pigmentosa (6), congenital stationary night blindness (CSNB) (7), diabetic retinopathy (8–10), and hypertensive retinopathy (11). Attenuated OPs are frequently observed in these conditions, sometimes preceding visible structural changes, supporting their potential as early biomarkers of retinal dysfunction (12).

Furthermore, OPs may serve as valuable functional biomarkers for assessing therapeutic efficacy in retinal interventions, including gene therapy. As OPs reflect integrated inner retinal function, they may capture downstream physiologic effects of treatment beyond the location of vector delivery or transgene expression (13). They may also aid in tracking retinal remodeling during disease progression and evaluating emerging therapeutic approaches such as optogenetics (14). Reliable and standardized quantification of OPs is therefore essential to advance diagnostic and therapeutic applications.

The standards for FFERG acquisition and reporting are well established and provided by the International Society for Clinical Electrophysiology of Vision (ISCEV). However, while these include a time-domain based description of OPs (i.e., amplitudes and latencies of individual wavelets), no consensus exists on standardized methods for the analysis of the frequency domain characteristics of OPs (15). OPs are challenging to characterize due to their transient nature and low amplitudes, making them susceptible to noise (Fig. 1A, left panel), hardware variability, and inconsistencies in recognition employing band-pass filtering (4, 16, 17). Most studies have focused on time-domain features of OPs, such as amplitude and peak timing (5, 18–20). However, these metrics may not fully capture the complexity of OPs, which are described here through additional frequency domain analyses (4, 21). Time-domain analysis examines how a signal changes over time, whereas frequency-domain analysis decomposes a signal into its constituent frequency components, offering insights into its spectral characteristics (17, 19, 20, 22). Relying solely on time-domain analysis risks overlooking critical OP features, making complementary frequency-domain analysis essential for a more comprehensive evaluation.

**Figure 1:**
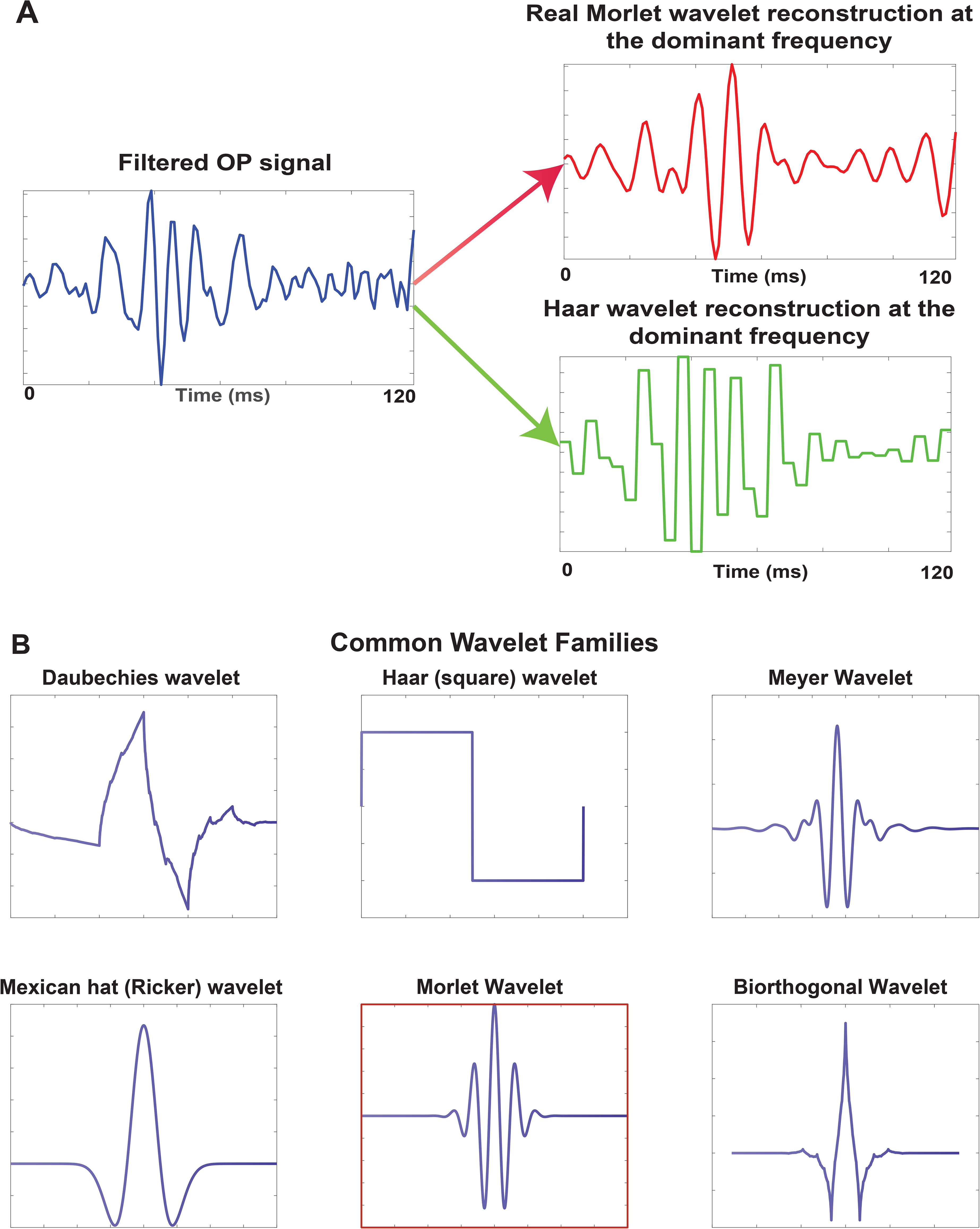
Wavelet families – method illustration. **A.** An example of an original noisy signal (on the left) following the application of a filter and its wavelet analysis using two different wavelets with decomposition (approximation) presenting a scale-specific wavelet component at a frequency close to the dominant frequency of the OPs in this example (∼ 100 Hz). The Haar wavelet approximation (bottom panel, green), previously unsuccessfully used in OP analyses (34), is compared to the Morlet wavelet approximation (top panel, in red), which was selected for this study as it closely resembles the original signal producing a better approximation as compared to the Haar wavelet. **B**. Several common wavelet families exist, and the success of signal analysis, such as OPs, depends on selecting the appropriate wavelet out of the wavelet families. The Morlet wavelet is highlighted in the red square.

Although the Fast Fourier Transform (FFT) is widely used to analyze the frequency-domain characteristics of time series (9, 18–20), it is poorly suited for transient signals like OPs. FFT assumes an infinitely long, stationary input signal, leading to artifacts and reduced accuracy when applied to short-duration signals like OPs (20). Alternative methods, including Short-Time Fourier Transform (STFT) and the Hilbert-Huang Transform (HHT), have been explored. STFT analyzes signals within fixed-width windows, improving time localization but sacrificing frequency resolution (21), which is particularly problematic for rapidly changing signals like OPs. HHT, a technique designed for nonstationary and nonlinear signals (23), shows promise in biological applications, but has not yet been widely adopted in OP analysis.

To overcome these limitations and enable comparison with existing methods, we investigated Continuous Wavelet Transform (CWT) analysis as a more suitable method for evaluating OPs (4, 16, 17). Unlike FFT and STFT, CWT does not assume signal stationarity and allows for simultaneous time and frequency localization, making it effective for detecting the brief, dynamic features of OPs (4, 17). We applied CWT alongside STFT-based approaches to FFERG analyses across experimental and clinical models of inner retinal dysfunction, including Royal College of Surgeons (RCS) and wild-type Brown Norway (BN) rats; *Atoh7* remote enhancer (REN) knockout (KO), *Atoh7* total enhancer knockout (TEN) KO, and wild-type mice; and human subjects with diabetic retinopathy and healthy controls. Our goal was to demonstrate that CWT provides a more physiologically meaningful and accurate assessment of inner retinal function in clinical and experimental settings.

## Methods

### Animal models

We utilized RCS rats (n = 10) as a model of retinal dystrophy (24). These animals possess a spontaneous null mutation in the Mertk gene, which disrupts RPE phagocytosis (24). In this rodent model, retinal degeneration begins around postnatal day 12 (P12) and results in the degeneration of almost all photoreceptors by P95, leading to blindness. Notably, a human counterpart of this model exists, leading to a Retinitis Pigmentosa phenotype (25). Healthy BN rats (n = 25) were used as a rodent model of intact retinal function, after ensuring the lack of advanced retinal disease previously reported in some of these animals by fundoscopy and electrophysiology (26).

Two genotypes of Atoh7 KO mice were used to assess inner retinal dysfunction resulting from disrupted retinal ganglion cell (RGC) development, as previously described (27). In Atoh7 REN KO mice (n = 3), the distal Atoh7 enhancer is deleted, leading to selective loss of RGCs, abnormal ipsilateral axonal projections, and partial optic nerve hypoplasia and retinal vascular abnormalities (27). In contrast, Atoh7 TEN KO mice (n = 6) carry a disruption of both proximal and distal enhancers, causing near-complete loss of RGCs and severe retinal vascular abnormalities (27). Since previous studies in human patients (12, 28) and animal models (29) have shown that retinal vascular abnormalities lead to attenuation of the OPs, we included the previously published Atoh7 models in this study (27). C57BL/6J wild-type mice (n = 6) were used as controls. We hypothesized that these genotypes exhibit graded deficits in inner retinal circuitry and synaptic signaling, spanning a range of dysfunction that impacts the dark-adapted FFERG OPs.

All procedures conformed to institutional and national ethical guidelines and were approved by the University of Pittsburgh Animal Care and Use Committee.

### Human subjects

We retrospectively analyzed FFERGs obtained from adults with diabetic retinopathy (n = 10, not stratified by severity, all non-proliferative disease) and healthy age-matched controls (n = 10), aged 26-67 years. ERG OPs are sensitive markers of inner retinal dysfunction and retinal circulatory abnormalities in diabetes, including patients without clinically apparent retinopathy (8, 10). Diabetes-related alterations in retinal vascular permeability and inner retinal signaling disrupt OP generation, resulting in characteristic changes in OP amplitude and timing (8, 10), providing a clinically relevant framework for comparing CWT to STFT. All procedures adhered to the Declaration of Helsinki and its subsequent amendments and were approved by the Institutional Review Board of the University of Pittsburgh (IRB #19030187).

### Data acquisition

For the rodent models, signals were sampled at 2000 Hz employing the Diagnosys Celeris system, while the human FFERG recordings were obtained at 1000 Hz sampling rate employing the Diagnosys Espion system (Diagnosys LLC, Lowell, MA). ERG waveforms were exported using Diagnosys eXporter software (Diagnosys LLC, Lowell, MA) into MATLAB-compatible data. All subsequent processing and analyses were performed in MATLAB R2026a (MathWorks Inc., Natick, MA). Standard ISCEV protocols were used (15) for both animals and humans, with proprietary Diagnosys Celeris electrodes, while human data were obtained employing Dawson, Trick, and Litzkow electrodes.

### Signal Preprocessing

To isolate the OPs from FFERG data, all waveforms were preprocessed using a 4-pole Butterworth high-pass filter with a cutoff frequency of 60 Hz in combination with a narrow-band 60 Hz notch filter to isolate the OPs and suppress electrical line noise, respectively. Signals were baseline corrected. Since the dominant frequency of dark-adapted oscillatory potentials tends to be lower in rodents than in adult humans, we used a 60 Hz high-pass cutoff instead of 75 Hz as accepted in clinical applications, accounting for lower characteristic OP frequency values. We maintained these filtering parameters for all data to increase uniformity.

### Short-Time Fourier Transform Analysis

To complement CWT, STFT was performed on filtered signals as described above. Power Spectral Density (PSD) and spectrogram plots were generated in MATLAB R2026a. An adaptation of the Discrete Fourier Transform (DFT), the STFT is used to obtain a time-frequency representation of signal power over time. In contrast to the DFT which describes the overall frequency content of a signal without providing adequate temporal localization, the STFT analyzes the signal within a sequence of finite analysis windows, allowing characterization of spectral changes as a function of time. In the STFT, the signal *x*(*t*) is multiplied by a finite-duration analysis window *w*(*t*–*τ*)centered at time τ, and the Fourier transform of the windowed segment is computed as 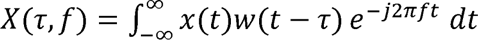. The spectrogram is defined as the squared magnitude of the STFT, *S*(*τ,f*) =| X(*τ,f*)|^2^, This allows for both temporal and frequency localization, making it more appropriate than DFT for analysis of nonstationary signals containing transient or evolving spectral features, such as the OPs.

Notably, because spectral estimates are derived from finite analysis windows, the temporal and spectral resolutions are interdependent, such that longer windows improve frequency resolution at the expense of time resolution and vice versa, a phenomenon termed “the uncertainty principle” (30). One-dimensional spectral density curves were derived from both STFT and CWT (31). For STFT analysis, the signal was segmented into overlapping windowed intervals, the Fourier transform was computed for each segment, and the squared magnitude of the resulting coefficients was normalized by sampling frequency and window energy to yield a segment-wise PSD. These local PSD estimates were then averaged across time to obtain a single Fourier-based PSD curve (nV²/Hz as a function of frequency). For all full-field ERG recordings a frequency resolution of 60 Hz was used. Although a finer resolution is achievable, the correspondingly longer analysis window spanned a large fraction of the approximately 150 ms oscillatory-potential epoch, truncating the recording’s edges and obscuring early OP components, whereas a 60 Hz resolution preserved the full analyzed epoch (see Fig. 2 for explanation of signal truncation). For the arbitrarily long synthetic validation signals, where this constraint does not apply, both a 10 Hz and a 60 Hz resolution were computed, the latter matching the real-data setting (Fig. 2).

**Figure 2:**
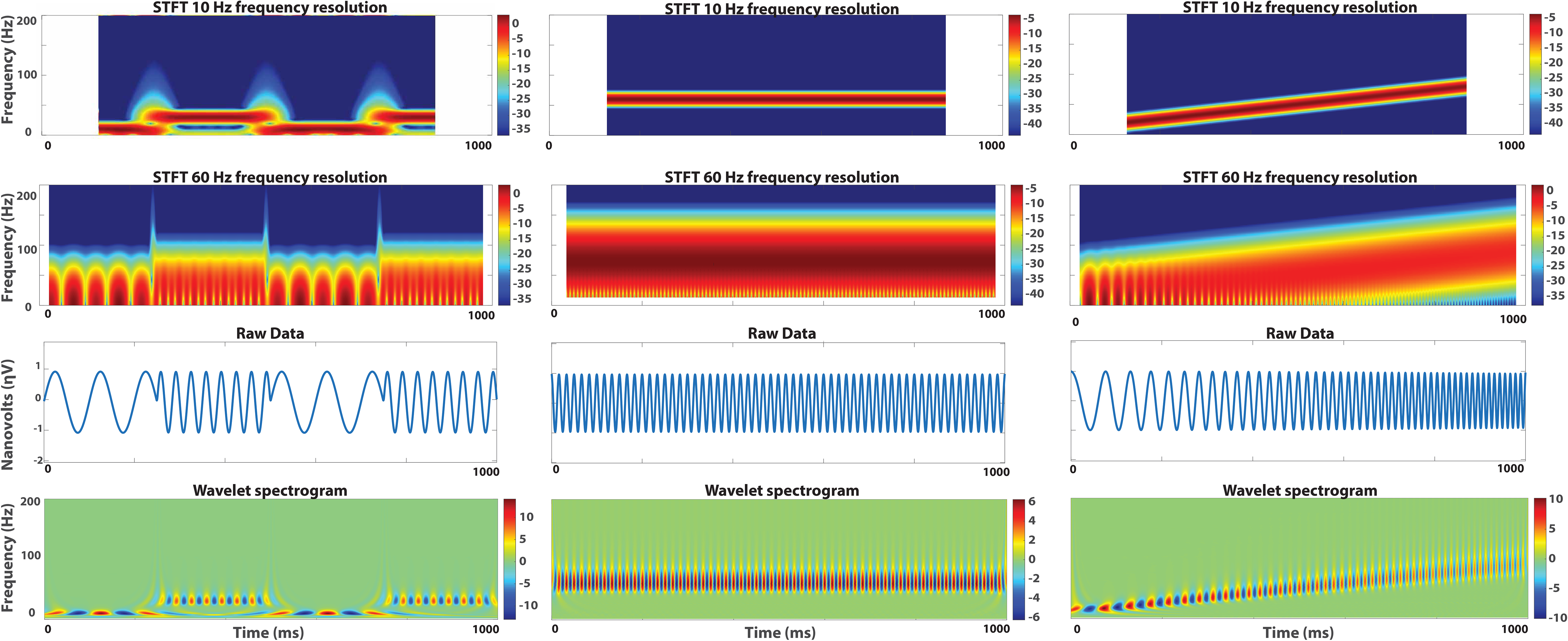
Analysis of unfiltered idealized signals: a 10–30 Hz alternating signal (left column), a 60 Hz sinusoid (middle column), and a 10–90 Hz chirp (right column). STFT with higher resolution (10 Hz, upper row) exhibited signal truncation for all signals and pronounced signal smearing in transition epochs (upper row, left panel). Reducing the resolution (60 Hz, second row) alleviated the truncation and smearing at the expense of poor frequency resolution, with difficulty recognizing the spectral composition of all signals, especially in the alternating signal (second row, left panel). CWT (lower row) spectrogram exhibited high fidelity of spectral analysis of the raw signal (third row), allowing accurate identification of the dominant frequencies for all signals as well as the temporal dynamics of the alternating signal (lower row, left panel).

### Wavelet Analysis

CWT decomposes a signal into localized time-frequency components using basis functions called wavelets. Two key parameters govern this process: (1) scale, inversely related to frequency, and (2) translation, determining the wavelet’s position in time. More formally, the CWT provides a representation of the signal across successive temporal scales. In contrast to Fourier-based methods, which decompose a signal into globally defined sinusoidal components, the CWT represents the signal using localized basis functions, or wavelets (Fig. 1B), generated by dilation and translation of a chosen wavelet basis function, or a “mother wavelet”, thereby permitting simultaneous temporal and scale localization of transient features. For a signal *x*(*t*), the CWT is defined as 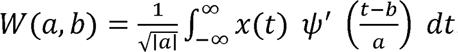, where *ψ* is the mother wavelet, *a* is the scale parameter, *b* is the translation parameter, and ’ denotes complex conjugation. The corresponding wavelet power may be expressed as *P*(*a*,*b*) =|*W*(*a*,*b*)|^2^. Herein, small scales emphasize higher-frequency, short-duration components, whereas larger scales emphasize lower-frequency, longer-duration structure. Because the transform is computed continuously across scales and positions, it provides a highly useful time-scale representation (32, 33). For wavelet analysis, the continuous wavelet transform was used to obtain time-scale coefficients. The squared magnitude of these coefficients was interpreted as wavelet power, and scale was converted to pseudo-frequency to permit representation on a frequency axis. A one-dimensional wavelet spectral density curve was then derived by averaging the wavelet power across time at each pseudo-frequency and applying a single multiplicative calibration constant to express the result as a spectral density in nV²/Hz. Because the continuous wavelet transform is a redundant (overcomplete) representation, the time-averaged wavelet power is proportional to the underlying spectral density but is not, on its own, correctly scaled in physical units; on a logarithmically spaced scale grid this constant of proportionality depends only on the mother wavelet and the sampling rate, and not on the analyzed signal. The calibration constant was therefore determined separately for each sampling rate by enforcing energy conservation (Parseval’s relation), such that the integral of the wavelet spectral density over frequency equaled the mean power (variance) of the analyzed signal. This was obtained from pure-tone reference signals of known power spanning the OP frequency band and was confirmed to be constant across that band. Thus, the STFT-derived curve represented a conventional Fourier PSD, whereas the wavelet-derived curve represented a CWT-based spectral density that allowed practical comparison of frequency-dependent power distributions, and specifically of the Peak Power Spectral Density (PPSD, the maximum value of the PSD curve) values, on a common frequency axis (31).

The real Morlet Wavelet was selected due to its resemblance to the morphology of OPs after filtering and its optimal balance between time and frequency localization (Fig. 1B). Compared to other wavelet families such as Daubechies and Haar, the Morlet Wavelet closely mirrored the oscillatory nature of OPs, preserving critical features while minimizing distortion with scale decomposition. In contrast, Haar, a frequently used wavelet family, produced a blockier, less physiologic signal approximation (Fig. 1A), as previously reported (34). The Morlet Wavelet thus offered reliable and biologically meaningful OP analysis.

Four visual outputs were generated. First, the raw ERG waveforms (microvolts as a function of time) were plotted to illustrate the unprocessed signal. Second, the filtered waveforms were plotted to visualize OP isolation. Third, spectrograms (frequency as a function of time) were generated using either CWT or STFT. Because the wavelet spectrogram displays signed wavelet coefficients on a linear scale whereas the STFT spectrogram displays power in decibels, the two spectrograms are shown on independent color scales and are not directly comparable in magnitude; they are presented to convey the temporal localization of frequency activity rather than absolute spectral power, the latter being quantified from the PSD curves. Finally, power spectral density (PSD) graphs were constructed, from which PPSD was derived for descriptive comparison using CWT/STFT ratios of group means. All preprocessing, frequency domain analysis using CWT and STFT and plots were performed and generated using MATLAB R2026a.

### Statistical Analysis

Dominant-frequency group values are reported as mean ± standard error of the mean (SEM), followed by units (Hz) and, for disease–control comparisons, the p-value. PPSD method comparisons are reported descriptively as CWT/STFT ratios of group means. For both CWT and STFT, OP spectral features were quantified using PSD plots and the dominant frequency (Hz) and Peak Power Spectral Density (PPSD, nV^2^/Hz) were derived from the filtered FFERG signals. For subjects or animals in whom recordings from both eyes were available, values from the two eyes were averaged to obtain a single subject-level value before group-level statistical analysis. CWT- and STFT-derived measurements obtained from the same recordings were compared descriptively, without hypothesis testing: PPSD as the dimensionless CWT/STFT ratio of group means (group mean CWT PPSD divided by group mean STFT PPSD), and dominant frequency as the mean within-recording difference (CWT − STFT) with its 95% confidence interval, calculated from the subject-level paired differences (t distribution, n − 1 degrees of freedom). Dominant-frequency comparisons between independent experimental or clinical groups were performed using Welch’s two-tailed t tests. Dominant frequency (DF) was defined as the frequency at which the spectral density reached its peak value (the PPSD) for each recording.

Group-level DF values were summarized using means and SEM; group mean PPSD and SEM are displayed descriptively in Figs. 3E, 4G and 5E, with PPSD method comparisons reported as CWT/STFT ratios of group means. Welch’s t tests were used for dominant-frequency disease-vs-control comparisons within each model: combined Atoh7 phenotypes (TEN KO and REN KO) vs. wild-type mice, RCS vs. BN rats, and diabetic retinopathy patients vs. healthy controls. All reported p-values are two-tailed. For the disease-vs-control DF endpoint within each model, the CWT- and STFT-derived Welch tests were treated as a family of two comparisons and Bonferroni-adjusted by multiplying the unadjusted p-values by 2 (maximum adjusted p = 1). Within-group comparisons of CWT-versus STFT-derived values from the same recordings were descriptive only; no hypothesis tests were performed on them, and no p-values are reported for them. Inferential analyses concern DF differences between independent experimental or clinical groups; within-recording method comparisons and all PPSD comparisons are descriptive. As expected from its superior frequency localization under equal-energy calibration, CWT consistently yielded higher and more spectrally concentrated peak values than STFT in the same recordings (Fig. 3E, 4G, 5E). Because this difference is, at least partially, expected from the frequency localization of the two transforms rather than being a hypothesis under test, it is reported descriptively as a CWT/STFT ratio of group mean PPSD, without hypothesis tests or p-values (see Limitations).

**Figure 3:**
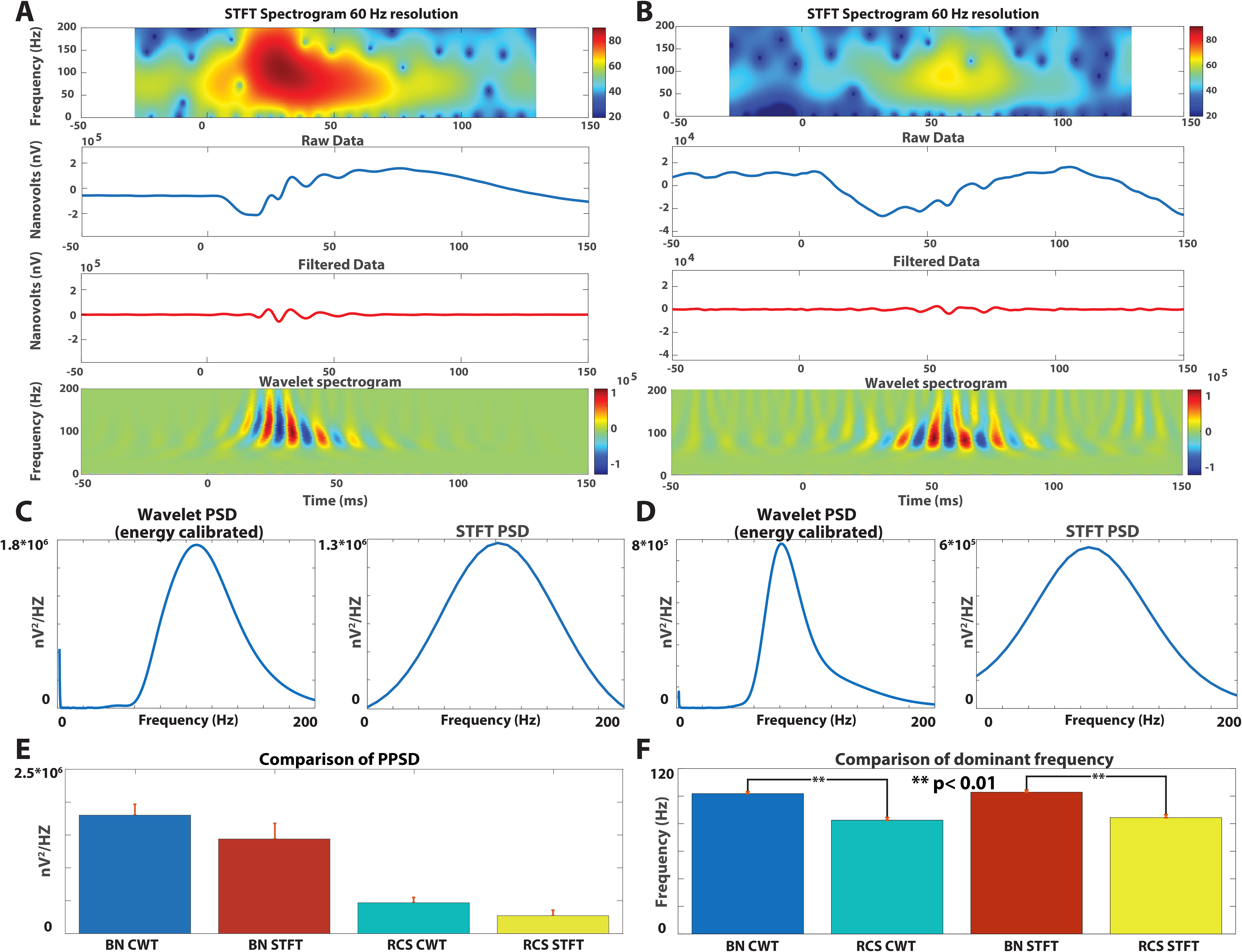
Comparison of CWT and STFT analyses in representative ERG recordings from control (panels **A**, **C**) and RCS (panels **B**, **D**) rats. **A** and **B** present the raw data (second row) and the filtered OPs (third row). STFT- (top row) and CWT-derived (bottom row) spectrograms are presented. Control ERGs exhibited prominent a- and b-waves (**A**, second row), with readily discernible oscillatory potentials (**A**, third row). RCS ERGs in this early stage (postnatal day 21) exhibited well-defined, morphologically preserved a-waves and markedly reduced b-waves (negative pattern, **B**, second row, note scale differences), while the OPs remained well defined (**B**, third row). Wavelet spectrograms demonstrate superior feature extraction (**A**, **B**, lower row), with the biological signal predominating the plot, while a considerable amount of noise is seen in the STFT spectrograms (**A**, **B**, upper rows). Wavelet PSD and STFT PSD are shown in **C** and **D** for the representative recordings in **A** and **B**, respectively. **E**. Descriptive group PPSD values (* 10 nV²/Hz). **F**. Group dominant frequencies (Hz). Bars show means and whiskers indicate SEM (BN, n = 25; RCS, n = 10). The CWT/STFT ratios of the group mean PPSD are 1.25 for BN and 1.74 for RCS rats (**E**), reported descriptively under equal-energy calibration without hypothesis tests or p-values. Brackets denote Welch’s BN–RCS comparisons within each method in **F** (CWT, Bonferroni-adjusted p = 4 × 10□¹□ ; STFT, Bonferroni-adjusted p = 4 × 10□□). **p < 0.01 for the indicated comparisons. Additional comparisons are reported in Results.

**Figure 4:**
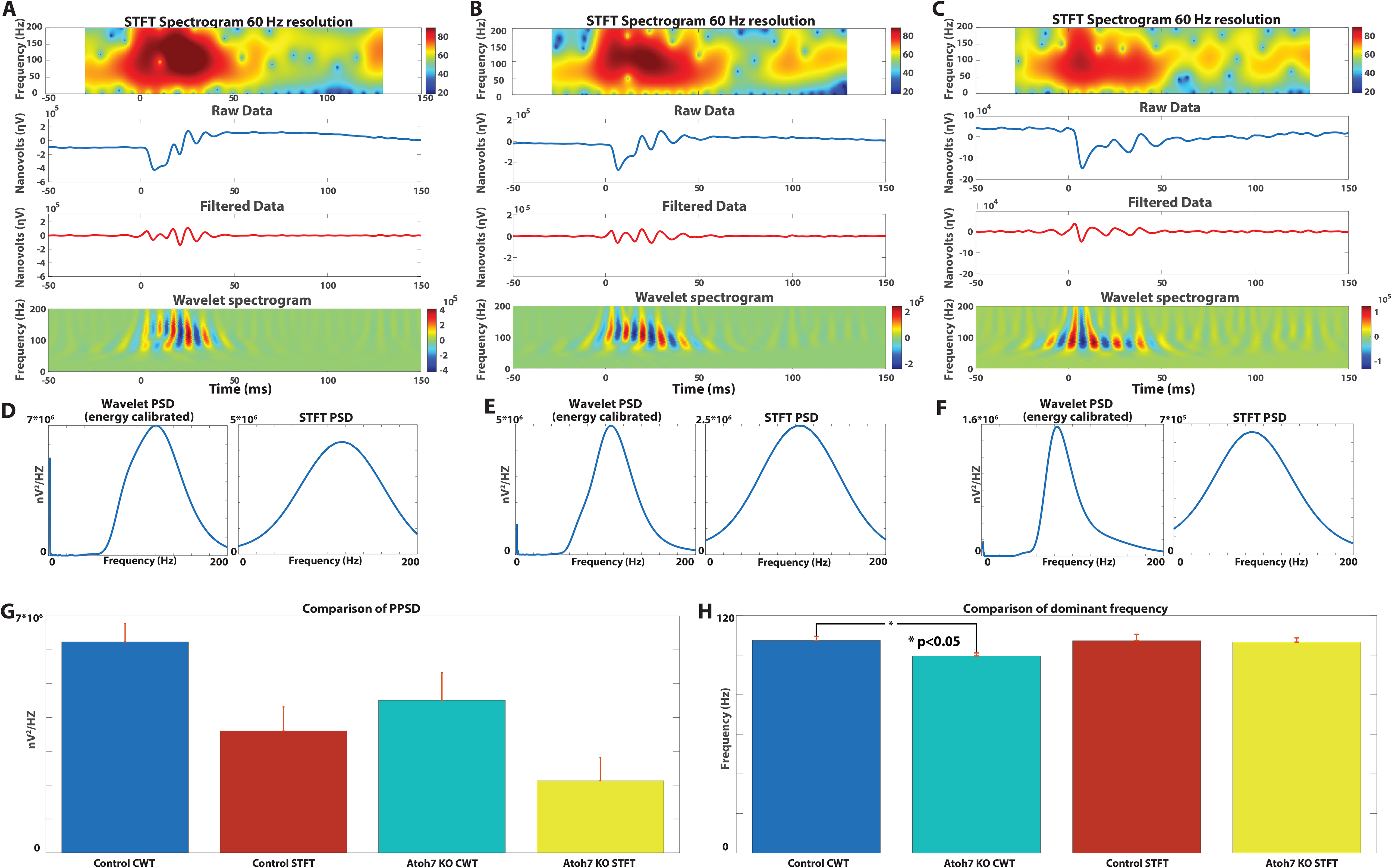
Comparison of CWT and STFT analyses in representative ERG recordings from control (panels **A**, **D**), Atoh7 REN (panels **B**, **E**), and Atoh7 TEN (panels **C**, **F**) mice. **A**-**C** present the raw data (second row) and the filtered OPs (third row). REN and TEN Atoh7 mice represented a spectrum of progressive dysfunction (**A**-**C**, second row), with lowering of amplitude in Atoh7 mice (**A**-**C**, second row) with a prominent negative pattern developing in Atoh7 mice (**C**, second row). STFT- (top row) and CWT-derived (bottom row) spectrograms are presented. Wavelet spectrograms again demonstrate superior feature extraction (**A**-**C**, lower row), with the biological signal predominating the plot, while a considerable amount of noise is seen in the STFT spectrograms (**A**-**C**, upper row). Wavelet PSD and STFT PSD are shown in **D**-**F** for the representative recordings in **A**-**C**, respectively. For statistical comparisons, the Atoh7 REN and TEN KOs were grouped together into a single Atoh7 KO group. **G**. Descriptive group PPSD values (* 10 nV²/Hz). **H**. Group dominant frequencies (Hz). Bars show means and whiskers indicate SEM (controls, n = 6; pooled Atoh7 KO, n = 9). The CWT/STFT ratios of group mean PPSD are 1.73 for controls and 2.11 for pooled Atoh7 KO mice (**G**), reported descriptively under equal-energy calibration without hypothesis tests or p-values. The bracket in **H** denotes the Welch’s control–KO comparison using CWT (Bonferroni-adjusted p = 0.027); *p < 0.05. The corresponding STFT comparison was not significant (Bonferroni-adjusted p = 1.00). Within KO mice, the CWT-derived dominant frequency was lower than the STFT-derived value (mean paired difference, CWT − STFT, −7.1 Hz; 95% CI [−9.8 to −4.4 Hz]; descriptive comparison). Other comparisons are reported in Results.

**Figure 5:**
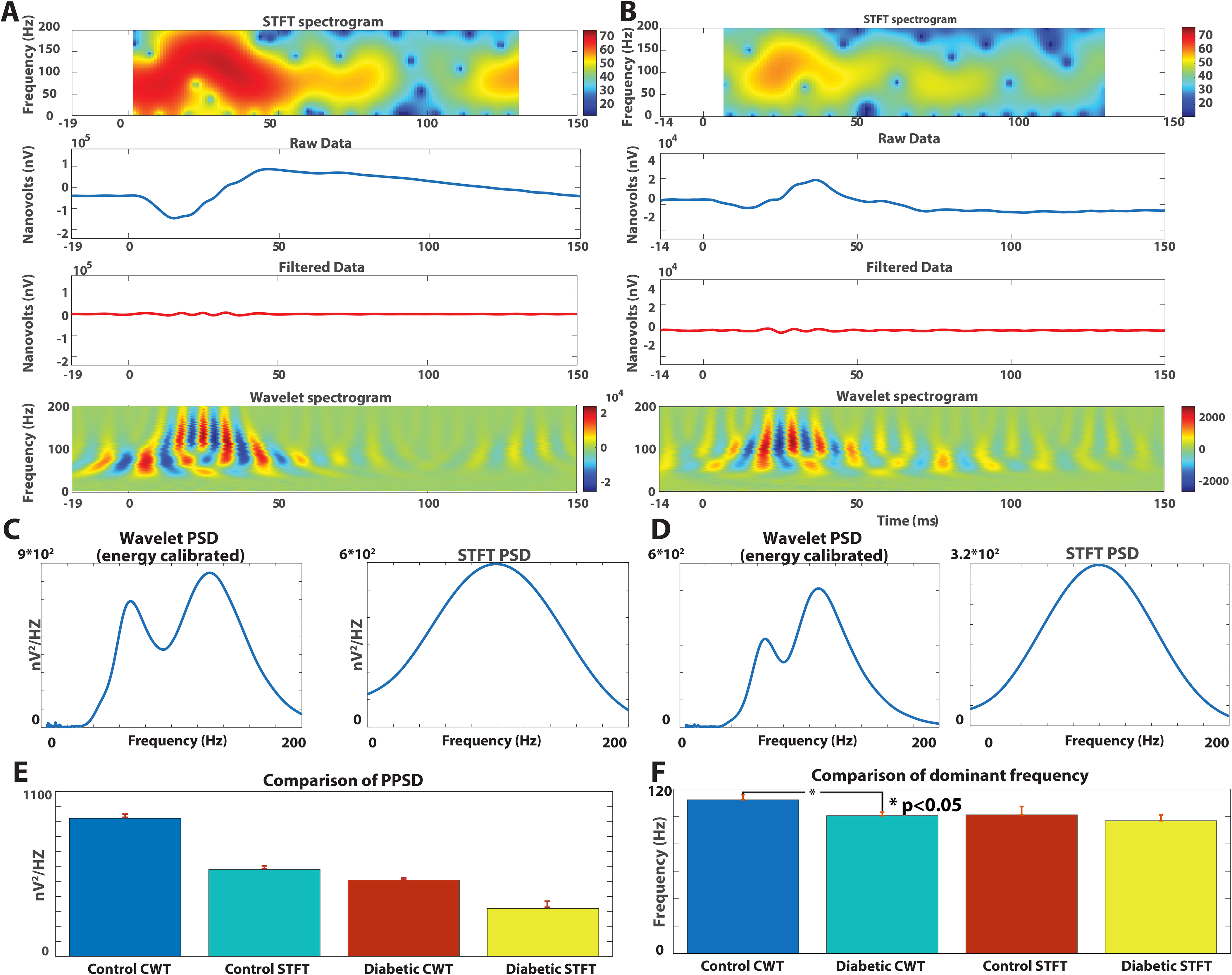
Comparison of CWT and STFT analyses in representative ERG recordings from control (panels **A**, **C**) and diabetic (panels **B**, **D**) patients. **A** and **B** present the raw data (second row) and the filtered OPs (third row). STFT- (top row) and CWT-derived (bottom row) spectrograms are presented. Control ERGs exhibited prominent a- and b-waves (**A**, second row), with readily discernible oscillatory potentials (**A**, third row), albeit these were of smaller amplitude relative to the a- and b-waves as compared to the rodent models (Figs. 3 and 4). In diabetic patients, the OPs were attenuated, with a markedly lower signal-to-noise ratio (**B**, third row), posing a challenging task for the analytical methods. Wavelet spectrograms demonstrate superior feature extraction (**A**, **B**, lower rows), with the biological signal predominating in the plot for control and the noise being clearly discernible in diabetic spectrograms utilizing CWT, while a considerable amount of noise is seen in the STFT spectrograms for both controls and diabetics (**A**, **B**, upper row). For diabetic patients specifically, the biological signal is not easily discernible from the background noise (**B**, upper row), limiting the visual separation of OP activity from noise in this representative recording. Wavelet PSD and STFT PSD are shown in **C** and **D** for the representative recordings in **A** and **B**, respectively. **E**. Descriptive group PPSD values (nV²/Hz) on a linear scale. **F**. Group dominant frequencies (Hz). Bars show the means and whiskers indicate SEM (n = 10 subjects per group). The CWT/STFT ratios of the group mean PPSD are 1.59 for controls and 1.60 for diabetic patients (**E**), reported descriptively under equal-energy calibration without hypothesis tests or p-values. The bracket in **F** denotes the Welch’s control– diabetic comparison using CWT (Bonferroni-adjusted p = 0.04); *p < 0.05. The STFT-derived between-group dominant-frequency difference was not significant after adjustment (Bonferroni-adjusted p = 1). Additional comparisons are reported in Results.

#### Validation with idealized signals

Given the nonstationary nature of OPs and known limitations of Fourier-based methods, additional validation was performed using synthetic signals with known frequency content. To validate the CWT and STFT implementations, both methods were applied to synthetic test signals with known frequency content, including a broadband chirp (10– 90 Hz), a signal alternating between 10 and 30 Hz and a pure sinusoidal signal at 60 Hz. The artificial signals were not pre-filtered. In addition to confirming that both transforms recovered the correct dominant frequencies, we verified that the calibrated wavelet and the STFT spectral densities each conserved signal energy: the integral of each spectral density over frequency matched the time-domain signal power to within a few percent (Parseval’s relation) (35). This ensured that Peak Power Spectral Density values derived from the two methods were expressed on a common, physically meaningful scale, so that any residual difference between them reflects time-frequency resolution rather than a difference in total power.

#### Validation with synthetic oscillatory potentials in recorded noise

To test frequency recovery under realistic conditions, synthetic OP-like bursts of known frequency were embedded in noise derived from the recordings themselves. For each cohort, the pre-stimulus (noise-only) segment of a representative recording was used to fit an eighth-order autoregressive model, from which surrogate noise of the full epoch length was generated and scaled to the measured pre-stimulus noise amplitude, preserving the spectral character and level of each cohort’s noise without periodic tiling artifacts. Bursts were modeled as Gaussian-windowed sinusoids (Gabor functions) with peak-to-peak amplitude matched to the filtered OP amplitude measured in the same cohort, so that the signal-to-noise ratio (SNR; burst RMS divided by noise RMS after filtering) was cohort-specific and ranged from 2.4 (diabetic patients) to 18.4 (BN rats). Three signal families were generated. (i) Single bursts (σ = 12 ms, centered 30 ms after stimulus onset) at 80–115 Hz in 5 Hz steps, with ten noise realizations per frequency per cohort (560 signals), to quantify dominant-frequency recovery as the absolute difference between the recovered and injected frequency. (ii) Paired bursts (σ = 6 ms) centered 25 and 40 ms after stimulus onset, i.e., 15 ms apart, with frequencies differing by 15 Hz (80/95, 90/105, 95/110 and 100/115 Hz), ten realizations per pair per cohort (280 signals), to test the resolution of closely spaced transient events. (iii) An SNR series at a fixed sampling rate of 1000 Hz, in which single bursts at 80–115 Hz were embedded in diabetic-patient noise with amplitude scaled to yield SNRs of 1, 2, 3.5, 6 and 10 (400 signals), to separate the effect of SNR from that of sampling rate. All synthetic signals were written in the native recording format at the sampling rate and epoch length of the source cohort and processed with the identical pipeline (filtering, transforms, spectral density estimation and dominant-frequency extraction) used for the biological recordings; the injected parameters were not available to the analysis, and the analyst was masked to the data content.

## Results

CWT and STFT analyses were applied to idealized data as well as OPs derived from FFERG across experimental animal models and human subjects, followed by synthetic signals. Across datasets, CWT provided time–frequency representations with reduced spectral dispersion compared with STFT (Figs. 2–5), supporting its utility for characterizing transient OP features not only in healthy animals, as previously suggested, but across different pathological phenotypes. In synthetic signals, CWT but not STFT were able to resolve closely-spaced frequency components, similar to what occurred in the human cohort.

### Idealized data

We first compared the performance of CWT and STFT on a set of artificial, or “idealized”, data for illustration of STFT limitations. Specifically, we used a 60 Hz sinusoid, a 10–90 Hz chirp and, a 10–30 Hz alternating signal without the pre-filtering used for biological signals. Results are summarized in Fig. 2, illustrating the difficulties employing STFT for transient signals. Improving the frequency resolution (10 vs 60 Hz resolution, with lower value representing better frequency segregation) required larger windows, truncating the signal edges and limiting the useful range of the spectrogram (Fig. 2, upper row). Alternatively, using lower frequency resolution alleviated the truncation of data from the edges of the spectrogram. However, using lower frequency resolution led to poor identification of discrete frequencies in the power spectrum analysis (Fig. 2, second row), with difficulty recognizing different frequencies in the alternating signal as the resolution approached the inter-frequency difference of the signal (Fig. 2, second row, left panel). When analyzing alternating signals (Fig. 2, left column), STFT spectrograms exhibited “transitional smear”, with little or no information present around the transition zones (Fig. 2, left column, upper two panels). No such issues occurred when employing wavelet analysis (Fig. 2, left column, lower panel), where the individual frequency epochs were readily discernible on the spectrogram. CWT accurately recovered the spectral composition across all test signals (Fig. 2, lower row), whereas STFT exhibited frequency smearing and reduced accuracy for nonstationary signals, struggling with “transition periods” (Fig. 2, upper 2 rows).

### RCS rat model

In BN controls the raw data exhibited a robust waveform, with well-defined a- and b-waves (Fig. 3A, second row) and readily discernible OPs (Fig. 3A, third row). In early stages of degeneration in RCS rats, the response gravitated towards a negative pattern (Fig. 3B, second row, note differences in axes compared to 3A), with well-defined and morphologically preserved a- and significantly reduced b-waves, as previously described. The OPs remained well defined in this model (Fig. 3B, third row). Given the high signal-to-noise ratio in both control and RCS animals, both STFT (Fig. 3A&B, top row) and CWT (Fig. 3A&B, bottom row) produced meaningful spectrograms, although the STFT spectrograms appeared to over-represent the noise component as compared to CWT spectrograms. Under equal-energy calibration, the CWT/STFT ratios of group mean PPSD were 1.25 for BN controls (n = 25) and 1.74 for RCS rats (n = 10; Fig. 3C–E). These ratios are reported descriptively, without hypothesis tests or p-values. CWT-based dominant frequency analysis revealed a significant difference between wild-type BN controls and RCS rats (BN 102.5 ± 1.0 vs RCS 83.3 ± 1.3 Hz, Bonferroni-adjusted p = 4 * 10^□10^; Fig. 3F). STFT analysis likewise detected a significant dominant-frequency difference between BN and RCS rats (103.8 ± 1.3 vs 85.0 ± 1.9 Hz; Bonferroni-adjusted p = 4 * 10^□7^; Fig. 3F). Within recordings, STFT- and CWT-derived mean dominant frequencies differed little in RCS rats (mean paired difference (CWT − STFT) −1.7 Hz, 95% CI [-4.3 to 0.9 Hz]; n = 10) and in BN rats (102.5 ± 1.0 vs 103.8 ± 1.3 Hz; mean paired difference −1.3 Hz, 95% CI [-2.8 to 0.2 Hz]; n = 25).

### Atoh7 mouse models

As in the rat model, the mouse controls exhibited a robust waveform, with well-defined a- and b-waves (Fig. 4A, second row) and readily discernible OPs (Fig. 4A, third row). The *Atoh7* REN KO showed attenuated responses, with a tendency towards lower a- and b-waves (Fig. 4B, second row) as well as lower OP amplitudes (Fig. 4B, third row). The frequency of the OP appeared lower in both STFT and CWT spectrograms (Fig. 4B, upper and lower panels, respectively). All the aforementioned changes appeared to be more pronounced in the *Atoh7* TEN KO (Fig. 4C), demonstrating a continuum of impairment between these models. In line with previous work describing inner retinal vasculature abnormalities in both *Atoh7* TEN KO and *Atoh7* REN KO, which in turn was shown to attenuate OPs in both animal models (29) and human patients (12, 28), were grouped together into a single group, termed *Atoh7* KO (n = 9). CWT analysis revealed a significant difference between the dominant frequencies of the control and KO groups (107.4 ± 2.1 vs 99.5 ± 1.6 Hz, Bonferroni-adjusted p = 0.027; Fig. 4H). The corresponding STFT-derived dominant-frequency comparison between the control and KO groups did not reach significance (107.3 ± 3.3 vs 106.6 ± 2.1 Hz, Bonferroni-adjusted p = 1.00; Fig. 4H). Within recordings, the mean CWT–STFT dominant-frequency difference was close to zero in controls (CWT − STFT, 0.1 Hz; 95% CI [-3.8 to 4.0 Hz]; n = 6), whereas CWT yielded a lower dominant frequency than STFT in Atoh7 KO mice (mean paired difference -7.1 Hz; 95% CI [-9.8 to -4.4Hz]; n = 9). The CWT/STFT ratios of group mean PPSD were 1.73 for control mice (n = 6) and 2.11 for pooled Atoh7 KO mice (n = 9; Fig. 4G), reported descriptively without hypothesis tests or p-values. In the representative recordings, STFT-based PSD curves exhibited broader spectral spread across frequencies (Fig. 4 D-F). Additionally, the STFT spectrograms showed reduced physiologic interpretability as compared to the CWT spectrograms, with background noise affecting the STFT spectrograms to a much larger extent (Fig. 4A-C, upper vs the lower rows), especially for the *Atoh7* TEN KO model in which the signal-to-noise ratio tended to be smaller as compared to control animals (Fig. 4C vs 4A).

### Human controls and diabetic retinopathy cohort

In human subjects, there was an obvious reduction in the amplitude of the OPs relative to the large a- and b-wave components of the dark-adapted scotopic response (Fig. 5 A&B, second panels, also see discussion). The responses in diabetic patients tended to be attenuated, with reduced OP amplitudes and a markedly worse signal to noise ratio. These were reflected in the spectrograms, with noise components predominating in the STFT spectrograms for both controls and diabetic patients, while the CWT spectrograms exhibited superior feature extraction in controls (Fig. 5A top vs bottom panels) and a clear separation of signal and noise components in diabetic patients (Fig. 5B top vs bottom panel). The CWT/STFT ratios of group mean PPSDs were 1.60 for diabetic patients and 1.59 for controls (n = 10 per group; Fig. 5E), reported descriptively without hypothesis tests or p-values. Mean dominant frequency was lower in diabetic patients than controls with both methods. The CWT-derived difference remained significant after Bonferroni adjustment (112.1 ± 3.7 vs 100.4 ± 2.7 Hz, adjusted p = 0.04; Fig. 5F). The STFT-derived difference was not significant after adjustment (101.0 ± 6.2 vs 96.8 ± 4.3 Hz, adjusted p = 1; Fig. 5F). Within recordings, the mean CWT − STFT difference in dominant frequency was 11.1 Hz in control subjects, with the CI excluding zero (95% CI [3.7 to 18.5 Hz]; n = 10) and 3.6 Hz in diabetics, with the CI including zero (95% CI [-2.1 to 9.3 Hz]; n = 10).

#### Validation with synthetic oscillatory potentials in recorded noise

Both methods recovered the frequency of single synthetic bursts embedded in each cohort’s noise across the entire 80–115 Hz range and the full range of SNR studied (Fig. 6A). Recovery error depended on the sampling rate rather than on SNR. In the 2000 Hz cohorts (rats and mice), the mean absolute error was 0.92–0.99 Hz for CWT and 1.95–2.49 Hz for STFT, a two-fold advantage for CWT that was maintained from the highest (BN, SNR 18.4) to the lowest (TEN, SNR 3.5) SNR (Fig. 6B). In the 1000 Hz human cohorts, the two methods were comparable (CWT 1.44–1.50 Hz, STFT 1.19–1.55 Hz). Because SNR and sampling rate were confounded across cohorts, SNR was varied independently at 1000 Hz (Fig. 6C, D). Recovery error was independent of SNR between 2 and 10 for both methods (CWT ≈ 1.4 Hz, STFT ≈ 1.2 Hz) and increased for both only at an SNR of 1; the values obtained in the real control and diabetic noise fell on these curves at their measured SNRs (Fig. 6C). The small, constant difference between methods at 1000 Hz reflected a systematic offset of the CWT estimate of approximately -0.65 Hz that was present at every SNR (Fig. 6D), whereas the descriptive spread of the two estimates was similar (SD ≈ 1.4 Hz for both). Recovery at physiological SNR was therefore limited by the frequency resolution of the analysis grids rather than by noise, and the frequency accuracy of the two methods at 1000 Hz was equivalent within the precision of the analysis.

**Figure 6:**
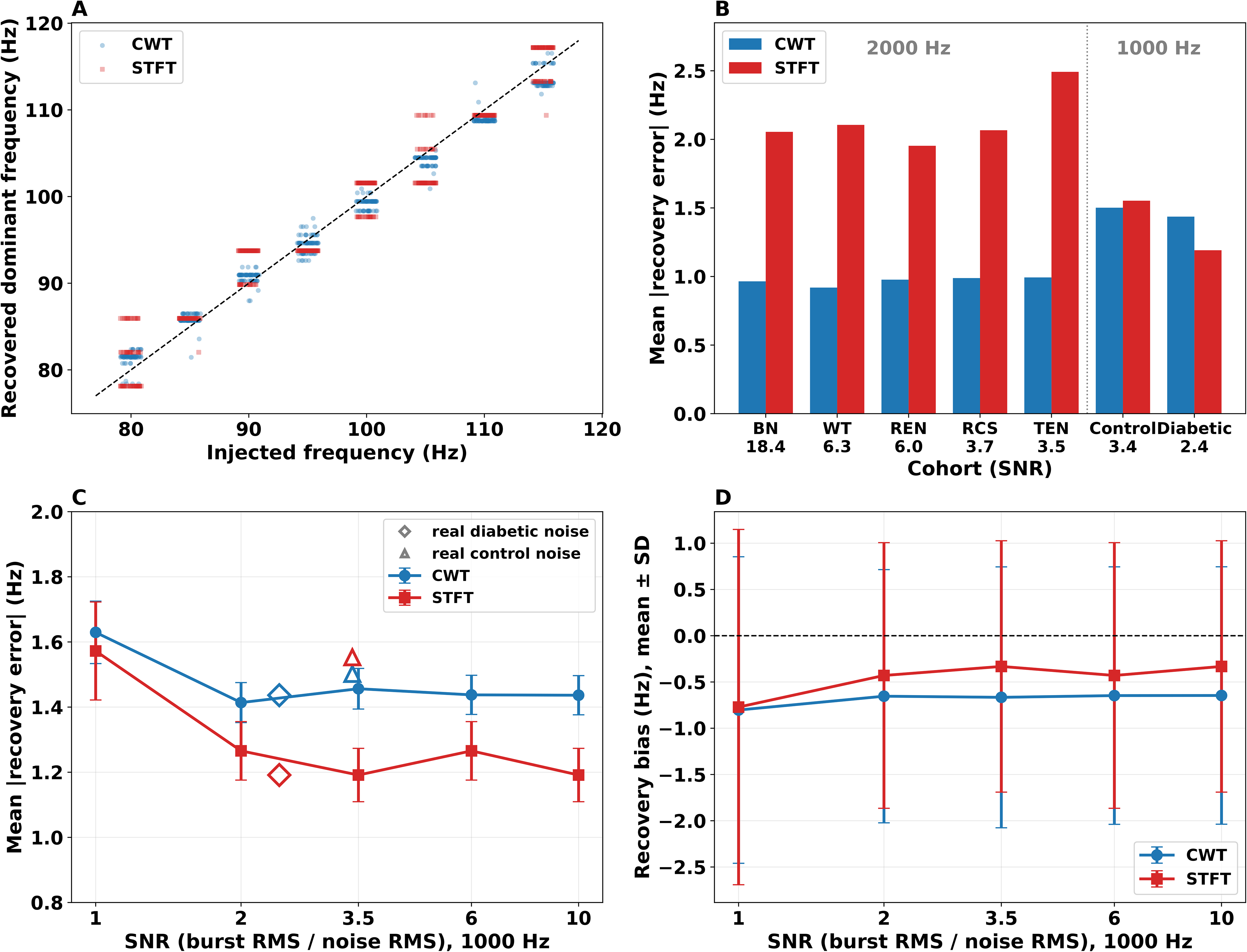
Validation of dominant-frequency recovery with synthetic oscillatory potentials embedded in recorded noise. Single Gaussian-windowed bursts of known frequency (80–115 Hz in 5 Hz steps; σ = 12 ms; centred 30 ms after stimulus onset) were embedded in autoregressive surrogates of each cohort’s own pre-stimulus noise, with amplitude matched to the cohort’s measured OP amplitude, and analyzed with the identical pipeline used for the biological recordings. **A**. Recovered versus injected dominant frequency for all 560 signals (7 cohorts * 8 frequencies * 10 noise realizations); dashed line, identity. **B**. Mean absolute recovery error per cohort, ordered from highest to lowest SNR; cohorts sampled at 2000 Hz (rats and mice) and at 1000 Hz (humans) are separated by the dotted line. At 2000 Hz, mean absolute errors were 0.92–0.99 Hz for CWT and 1.95–2.49 Hz for STFT; at 1000 Hz, they were 1.44–1.50 Hz and 1.19–1.55 Hz, respectively. **C**. Mean absolute recovery error (± SEM) as a function of SNR at a fixed sampling rate of 1000 Hz (diabetic-patient noise; burst amplitude scaled to SNR 1–10; 80 signals per level). Open symbols show the values obtained in the real control (triangle) and diabetic (diamond) noise at their measured SNR. **D**. Recovery bias (mean ± SD) for the same signals; dashed line, zero bias. The CWT estimate showed an approximately −0.65 Hz systematic offset across SNR levels, with similar spread for both methods (SD ≈ 1.4 Hz). CWT, blue; STFT, red.

Importantly, the two methods diverged when two bursts of different frequency were placed 15 ms apart (Fig. 7). The wavelet spectrogram resolved the two bursts as temporally distinct oscillatory events in every cohort, including the diabetic noise at the lowest SNR (Fig. 7B, E), whereas the STFT spectrogram merged them into a single, structureless region spanning most of the OP band (Fig. 7C, F). In contrast, the dominant frequency of the time-averaged spectral density failed to report either injected frequency for both methods (Fig. 7G–I). The CWT-derived peak occurred at an intermediate frequency determined by spectral interference between the temporally overlapping bursts and was nearly identical across all seven cohorts for a given pair (e.g., 85.3–86.2 Hz for the 100 + 115 Hz pair), indicating a property of the signal rather than of the noise. The STFT-derived peak was confined to a single frequency bin (93.8 Hz) for all four pairs in the human cohorts. Time-averaged spectra from either method were thus unable to characterize closely spaced transient events, which were resolved only by the time–frequency representation of the wavelet transform.

**Figure 7:**
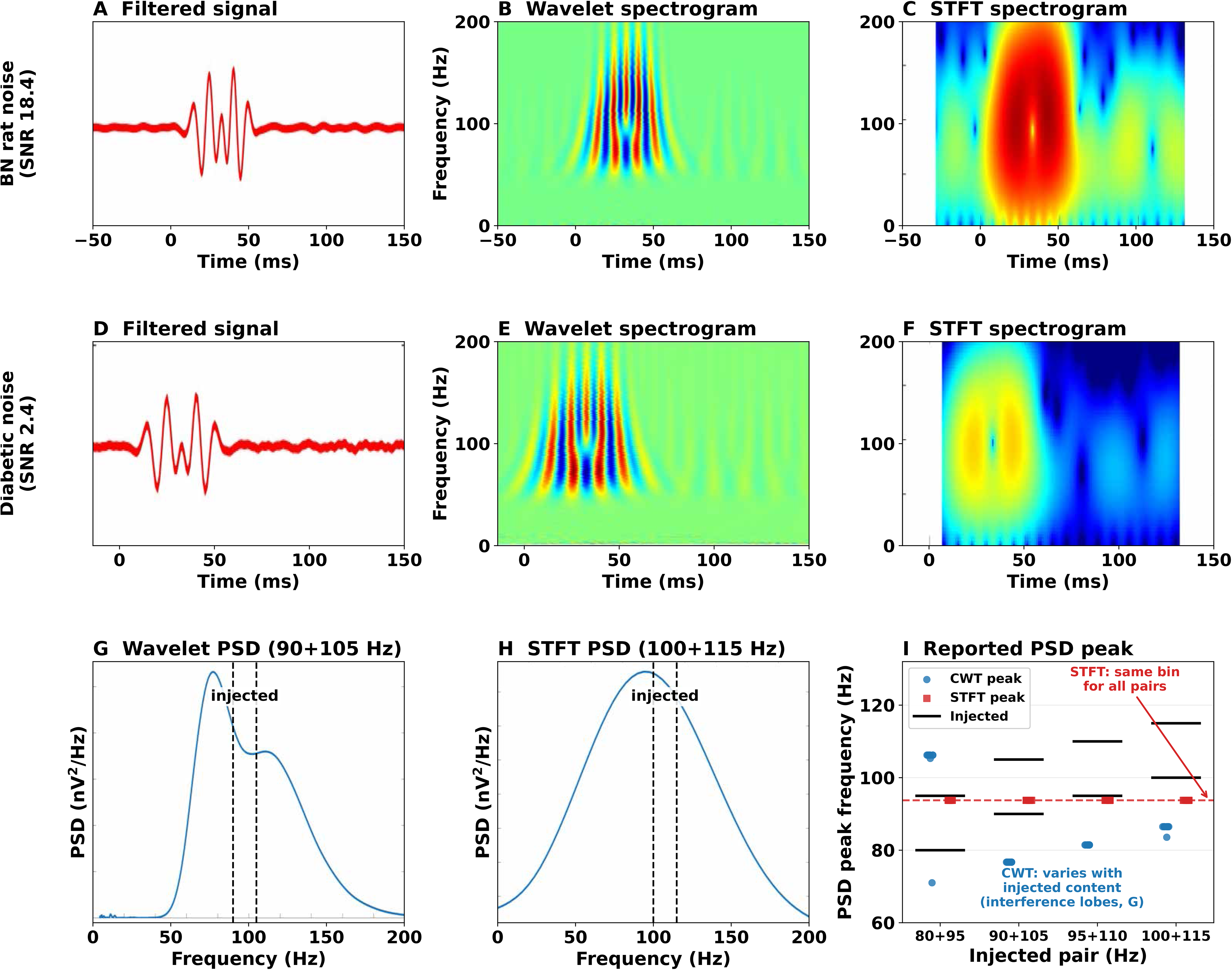
Only wavelet analysis resolves closely spaced oscillatory events under the evaluated settings. Two Gaussian-windowed bursts (σ = 6 ms) separated by 15 ms (90 Hz at 25 ms and 105 Hz at 40 ms) were embedded in autoregressive surrogates of recorded pre-stimulus noise with cohort-matched amplitude. **A–C.** High-SNR condition (BN rat noise, SNR 18.4): the filtered signal (**A**) contains both bursts; the wavelet spectrogram (**B**) resolves them as temporally distinct oscillatory events, whereas the STFT spectrogram (**C**) merges them into a single unresolved region that reflects the envelope of the oscillatory epoch rather than its individual components, with energy broadly dispersed along the frequency axis. White margins in **C** and **F** reflect the edge truncation imposed by the STFT analysis window. **D–F.** The same result at the lowest cohort-matched SNR (diabetic-patient noise, SNR 2.4). The shallow amplitude modulation discernible in the STFT (**F**) reflects the envelope minimum between the two bursts, not their resolution: both lobes occupy the same broad frequency band, and neither component’s frequency can be assigned. **G, H.** Time-averaged spectral densities (dashed lines, injected frequencies): the wavelet PSD of the recording shown in **D–F** (**G**) exhibits interference structure with two identifiable peaks displaced from both injected frequencies, whereas the STFT PSD, shown for a 100 + 115 Hz pair in the same noise (**H**), forms a single broad envelope whose maximum (93.8 Hz) lies below both injected frequencies and is essentially identical for every pair (**I**). **I.** Dominant frequency of the time-averaged spectral density for all four injected frequency pairs (10 noise realizations each, diabetic noise): neither the wavelet- nor the STFT-derived peak reports either injected frequency (black bars). The wavelet-derived peak corresponds to the taller interference lobe (**G**) and varies with the injected pair, whereas the STFT-derived peak is identical for all pairs (dashed line, 93.8 Hz), insensitive to the injected content. Time-averaged dominant frequencies from either method therefore failed to recover the injected components, whereas the wavelet spectrograms resolved the events in time (**B**, **E**). Spectrograms and spectral densities use their native, independent scales (signed wavelet coefficients; STFT power in dB for spectrograms); the comparison concerns time–frequency structure, not magnitude.

## Discussion

In this study, we evaluated CWT and STFT approaches for characterizing the OPs of the dark-adapted FFERG across experimental and clinical models of inner retinal dysfunction. Our findings demonstrate that CWT, particularly using the real Morlet wavelet, provided a physiologically grounded representation of OP time–frequency structure across the models examined. In our analyses, CWT consistently reduced spectral dispersion, preserved temporal localization on the spectrogram through superior spectral feature extraction, and provided visually clearer representations of transient OP activity than STFT, including in attenuated signals from disease models and patients. Here, spectral dispersion refers to the spread of signal content across frequencies within a recording, not to between-subject variance in extracted PPSD or dominant-frequency values.

OPs are believed to reflect the activity of the inner retinal circuitry arising from interactions among bipolar cells, amacrine cells, and retinal ganglion cells (5, 36). Their sensitivity to inner retinal dysfunction in human patients has been documented in retinitis pigmentosa, CSNB, diabetic retinopathy, and hypertensive retinopathy (6–8, 10, 11). However, the transient, high-frequency nature of OPs poses significant challenges for traditional frequency-domain analysis, which often assumes signal stationarity and long-duration oscillations (9). Consistent with prior work, our results reinforce the understanding that OPs are fundamentally nonstationary signals characterized by rapid temporal evolution and frequency modulation (4, 17). Unlike FFT- or STFT-based approaches, CWT does not assume stationarity and instead adapts temporal and frequency resolution dynamically, making it well suited to capture the brief oscillatory bursts that define OPs (16, 20).

There are several theoretical and practical reasons for the better performance of CWT as compared to STFT when analyzing OPs. FFT, the classical method of frequency analysis of stationary signals (5, 9, 18–20), does not provide adequate time resolution in the frequency domain for transient signals like the OPs. It assumes a stationary and infinite signal, introducing various artifacts when these assumptions are violated, as is the case with OPs. Notable artifacts, discussed below, include spectral leakage, broadband spreading, and ringing/Gibbs phenomenon (37), in addition to the poor temporal localization mentioned above (38, 39). STFT seeks to overcome these limitations by dividing signals into limited time windows, leading to improved time localization, within which the classical FFT is performed. However, this comes at the expense of frequency resolution, as shorter time windows impose natural frequency analysis limitations for FFT (21). This phenomenon of frequency-time resolution tradeoff, termed “the uncertainty principle”, has been described in classic works of signal processing, showing that the lower bound of the time-bandwidth product for an analysis window is (4π)^−1^ (33). Notably, each window is still subject to the same artifacts mentioned above. Spectral leakage occurs when energy from a finite signal segment spreads into adjacent frequency bins, causing narrowband components to appear broader than their true spectral extent, as can be seen from the broad distribution of the spectral power of the OPs analyzed by STFT, more so in noisy signals (e.g., Fig. 4C and F for *Atoh7* TEN KO). Broadband spreading, seen at the transition periods in the alternating-frequency sinusoid in our idealized data (Fig. 2, left column), is common for transient events, whose short duration inherently distributes energy across a wide frequency range. This occurs since STFT (or essentially FFT performed in each window) collapses the entire analyzed segment into a single global spectrum without preserving the timing of individual events, which appears as diffuse high-frequency activity. Windowing also leads to spectrogram “clipping” or truncation, where enlarging the analysis window to achieve better frequency resolution artificially shortens the analyzed signal (Fig. 2, upper vs second row). This occurs because, with centered windows, the first and last valid spectrogram times lie approximately one-half window length inside the signal boundaries, leading to the apparent shortening of the time axis, worsening as larger windows are used (31). Artificially extending the signal is a method used to overcome such issues, which can be achieved by zero padding or duplication. However, these create abrupt signal transitions prone to “ringing”, or Gibbs phenomenon artifact, where artificial oscillatory activity appears because of an abrupt transition, such as those seen in artificially extended signals by zero padding (Supplementary figure 1).

Notably, the uncertainty principle applies to the wavelet transform as well, albeit in a more complicated manner. In wavelet analysis, the mother wavelet is dilated and translated to fit the signal of interest, changing the effective window at each scale. At small scales the wavelet is compressed, yielding better time localization but poorer frequency localization; at large scales the wavelet is stretched, yielding better frequency localization but poorer time localization. Thus, wavelet analysis offers adaptive or scale/frequency dependent resolution, with finer time resolution at high frequencies and finer frequency resolution at low frequencies (32). In practice this is much better suited for the analysis of biological signals, making CWT more appropriate for this aim as compared to STFT. However, the uncertainty principle limiting the lower bound of the time-bandwidth product for each scale to (4π)^−1^ still applies (33).

The choice of wavelets has a profound effect on the success of wavelet analysis. The real Morlet wavelet used in this study closely matched the morphology of filtered OPs, enabling faithful reconstruction of oscillatory structure while minimizing spectral distortion, in agreement with previous wavelet-based analyses (Fig. 1A) (4, 17). This occurs since the wavelet transform coefficients are derived from correlations between the signal and the scaled and time-shifted versions of the selected wavelet (32). Thus, choosing a “mother wavelet” function bearing close resemblance to the analyzed signal improves wavelet analysis by means of better signal reconstruction fidelity and efficiency (Fig. 1A) (40). Employing wavelets of similar morphology to the analyzed signals leads to larger and more concentrated coefficients improving localization and feature extraction (Figs. 3-5). This principle is relevant in many biomedical signals and is not necessarily limited to OPs. It has been shown in other medical specialties that transient waveform shape often carries diagnostic significance, and signal-dependent wavelet selection can improve compression and analytic performance (41, 42).

Across sample data and experimental models examined in this study, CWT spectrograms provided clearer temporal localization and physiologic interpretability of transient OP activity than STFT spectrograms. In this study we focused on models known to have an inner retinal deficit.

Although STFT improves upon conventional FFT by introducing temporal localization of the spectrogram spectral properties, the uncertainty principle resulted in spectral smearing and reduced frequency specificity (19, 21) (Figs. 2-5), consistent with prior reports cautioning against fixed-window Fourier methods for transient neural oscillations (16, 18, 20). These limitations were further supported by validation analyses using synthetic (“idealized”) signals with a known spectral content, in which CWT accurately recovered expected dominant frequencies while STFT demonstrated reduced accuracy for nonstationary inputs (Fig. 2). Notably, the spectral smear worsened with higher resolution as use of larger windows resulted in the transition zone being present in multiple analysis windows (Fig. 2, left column, upper panel for higher and second panel for lower resolution).

In BN controls and RCS rats, both CWT and STFT produced interpretable results due to the relatively high quality of the ERG recordings (Fig. 3). Both methods detected disease-related alterations in OP frequency content, consistent with previously described secondary inner retinal remodeling following photoreceptor degeneration (43). PPSD comparisons are summarized descriptively by CWT/STFT ratios of group means of 1.25 in BN controls and 1.74 in RCS rats, consistent with greater spectral concentration under equal-energy calibration. Both methods also detected significantly lower dominant frequencies in RCS rats (CWT, Bonferroni-adjusted p = 4 × 10□¹□; STFT, Bonferroni-adjusted p = 4 × 10□□). Within recordings, CWT- and STFT-derived mean dominant frequencies differed little in either rat cohort (mean paired differences, CWT − STFT: RCS, −1.7 Hz, 95% CI [−4.3 to 0.9 Hz]; BN, −1.3 Hz, 95% CI [−2.8 to 0.2 Hz]). These findings support the use of both transforms to characterize degeneration-related OP alterations in this cohort.

In both the *Atoh7* REN and TEN KO mouse models, an impairment in the inner retinal blood supply has been described (27), which in turn is known to cause attenuation of the OPs (44). The electrophysiological correlate of this dysfunction is clearly seen when looking at the ERG traces (Fig. 4A-C, second row). The substantial noise component in the highly attenuated recording from the *Atoh7* TEN mouse (Fig. 4C) diminished the usefulness of the STFT-spectrogram (upper panel), while the CWT spectrogram exhibited a clearer separation between the biologically relevant signal and noise components (lower panel). Pooling the *Atoh7* REN and TEN KO mice into a single group and comparing them to control is justified by the common pathophysiology of inner retinal damage in these models, stemming from abnormal inner retinal blood supply (27). The CWT-based dominant-frequency comparison remained significant after Bonferroni adjustment, whereas the STFT-based comparison did not (Bonferroni-adjusted p = 0.027 for CWT and 1.00 for STFT; Fig. 4H). PPSD comparisons are summarized descriptively by CWT/STFT ratios of group means of 1.73 in control mice and 2.11 in pooled Atoh7 KO mice (Fig. 4G), without hypothesis tests or p-values. Within recordings, CWT-derived dominant frequency was lower than STFT-derived dominant frequency in the KO group (mean paired difference, CWT − STFT, −7.1 Hz; 95% CI [−9.8 to −4.4 Hz]), whereas the mean paired difference in controls was 0.1 Hz (95% CI [−3.8 to 4.0 Hz]). These within-recording comparisons are descriptive. While we are currently obtaining more recordings to improve our sample size for each sub-group, these method-dependent dominant-frequency findings warrant confirmation in larger cohorts, particularly given the limited sample size (45). However, our analysis points towards greater biological interpretability of the CWT-based spectrograms, with CWT (but not STFT) based dominant frequency differences reaching statistical significance. The present results therefore support further evaluation of CWT-based OP metrics in these models.

In human subjects with diabetic retinopathy, an additional analytical difficulty was introduced. Relative to total ERG amplitudes, rodent OPs are more prominent and have larger relative amplitudes as compared to humans (46). This stems from smaller a-waves and slower b-waves in the rodents, making the OP relative amplitude appear larger (44), improving the yield of all analytical methods (47). The noise component was readily observable in control (Fig. 5A) and diabetic (Fig. 5B) patients. For control patients, the noise component appeared to be dominant throughout the STFT-spectrogram, while in the wavelet spectrogram the ability of the analytical method to accentuate the biological signal effectively overcame this limitation (Fig. 5A top vs bottom row). In the setting of the more challenging diabetic ERG with attenuated signal and substantial noise contamination, the STFT spectrogram provided almost no means to discern meaningful signal from background noise (Fig. 5B, upper row). The wavelet spectrogram, while displaying a significant noise signal, created a clear separation between the background noise and the biological signal, with the latter being of visibly larger intensity (Fig. 5B, lower panel). This separation carried into the one-dimensional spectral summaries: because the time-averaged PSD weights each component by its duration rather than its instantaneous intensity, the temporally persistent noise formed a distinct lower-frequency peak alongside the dominant OP peak in the wavelet PSDs of both groups (Fig. 5C, D). The STFT, who’s fixed 60 Hz frequency resolution is coarser than the spectral separation between the two components, instead merged them into a single broad peak whose maximum fell between them — at a frequency corresponding to neither the OPs nor the noise — recapitulating in clinical recordings the behavior demonstrated with synthetic two-component signals (Fig. 7G–I). The descriptive CWT/STFT ratios of group mean PPSD were 1.60 in diabetic patients and 1.59 in controls (Fig. 5E); no PPSD hypothesis tests or p-values are reported. Both methods also showed a lower mean dominant frequency in diabetic patients than controls. The CWT-derived comparison remained significant after Bonferroni adjustment (controls, 112.1 ± 3.7 Hz; diabetic patients, 100.4 ± 2.7 Hz; adjusted p = 0.04), whereas the STFT-derived comparison did not (controls, 101.0 ± 6.2 Hz; diabetic patients, 96.8 ± 4.3 Hz; adjusted p = 1; Fig. 5F). Within recordings, the mean CWT − STFT difference in dominant frequency was 3.6 Hz in diabetic patients (95% CI [−2.1 to 9.3 Hz]) and 11.1 Hz in controls (95% CI [3.7 to 18.5 Hz]). These paired differences are reported descriptively. These separate disease–control comparisons do not constitute a direct test of differences in diagnostic performance between methods. The above observations align with prior studies linking OP abnormalities to early inner retinal dysfunction and impaired neurovascular coupling in diabetes (10, 12) and support the utility of CWT in the analysis of ERGs obtained in a clinical setting. There, patient comfort considerations (e.g., the use of skin vs contact lens electrodes) and inherent signal differences as compared to rodent models (44, 46) lead to smaller signal-to-noise ratios, requiring more sensitive analytical methods to accurately discern the biological signal from background noise (48). In particular, skin electrodes placed near the lower eyelid record substantial periorbital (orbicularis oculi) muscle activity, whose surface-EMG power spans approximately 20–500 Hz and therefore overlaps the OP band entirely, rendering this noise refractory to frequency-domain filtering (49, 50). To summarize, only CWT (and not STFT) was able to resolve separate frequency components within the signal, leading to a significant difference in the reported dominant frequency. These results point towards an advantage of employing CWT in the context of noisy biological signals, such as the OPs, in clinical settings.

To better explore the apparent advantage of employing CWT in the setting of separate frequency components within the signal, we performed validation with synthetic OP-like bursts embedded in the recorded noise of each cohort extended these observations to large-scale datasets under realistic conditions (Figs. 6 and 7). Both methods recovered the frequency of an isolated burst to within 1–2.5 Hz across the full range of physiological SNR, confirming the validity of the analysis pipeline and of the energy calibration; CWT showed approximately half the mean absolute frequency-recovery error at 2000 Hz, whereas both methods showed similar error at 1000 Hz, where a small (≈0.65 Hz), SNR-independent systematic offset of the wavelet estimate accounted for the residual difference That recovery error did not vary with SNR between 2 and 10 indicates that, at physiological SNR, dominant-frequency precision is set by the resolution of the analysis grid rather than by noise. The decisive difference emerged for two bursts separated by 15 ms: only the wavelet spectrogram resolved the individual events, the STFT merged them, and the dominant frequency of the time-averaged spectrum — from either method — reported neither component. This last result is of practical importance for 2 main reasons. (1) OPs are a train of closely spaced wavelets, and a single dominant frequency extracted from a time-averaged spectrum is an inherently limited descriptor of such a signal irrespective of the transform used. The information distinguishing the individual OP components resides in the time–frequency plane, where the CWT showed a clear advantage in resolving closely-located frequency components. Future applications using CTW could, potentially, employ reporting strategies beyond the single frequency reporting as described here, aimed at better characterization of the frequency composition of the OPs. (2) Furthermore, as demonstrated by the results of the human cohort above and validated by synthetic signals, CWT offers a distinctive advantage in the setting of noisy biological signals, where the noise component’s frequency composition is similar to that of the biological signal.

Despite well-established ISCEV standards for ERG acquisition and reporting (15), no consensus exists regarding standardized methods for frequency analysis of OPs. Most clinical and experimental studies continue to rely on time-domain metrics or Fourier-based spectral analysis, which may obscure clinically meaningful information embedded in OP dynamics (8, 9, 19, 51). Our findings suggest that CWT offers a path toward more reliable and physiologically grounded OP quantification, improving signal spectral feature extraction by preserving the temporal dynamics while providing high-resolution spectral information. Its multiresolution structure allows better temporal localization of high-frequency transient events while preserving better frequency localization of lower-frequency components (52). In contrast, the fixed window of the STFT imposes a uniform time-frequency tradeoff across the entire spectrum (the uncertainty principle), which may reduce sensitivity to short-duration or multiscale features (31). This property makes wavelet methods particularly advantageous for biomedical signals containing transient, evolving, or morphologically complex patterns, such as OPs. Feature extraction is also improved through wavelet analysis and spectrograms for noisy signals, as is the case with OPs in both clinical and research settings (Figs. 1A, 3-5). Transient signal components tend to be concentrated into a relatively small number of wavelet coefficients, whereas background noise is often distributed more diffusely across scales and positions. This difference can improve the contrast between relevant signal features and noise, thereby facilitating denoising and preservation of diagnostically meaningful waveform structure during reconstruction aimed at feature extraction in subsequent analyses, not explored in this paper (32). In contrast, STFT uses a fixed analysis window, which can spread short-duration features over broader time-frequency regions and reduce their separation from background noise, particularly when the signal is nonstationary (53).

CWT-based OP analysis may therefore enhance the translational utility of ERGs in both research and clinical settings. Improved characterization of inner retinal function has important implications for diseases in which OPs serve as early biomarkers, including diabetic retinopathy, inherited retinal dystrophies, and disorders of retinal circulation (6, 9, 10, 54). Moreover, CWT-based analysis may prove particularly useful in assessing the efficacy of emerging therapies affecting inner retinal function, such as intravitreal gene therapy, neuroprotective approaches, and optogenetic interventions (14).

Several limitations should be acknowledged. Sample sizes were small in certain experimental groups, particularly *Atoh7* KO mice, which limits the precision of group estimates and the ability to detect differences using either method. The reported dominant-frequency p-values describe individual comparisons and do not constitute a direct test of differences in statistical power or diagnostic performance between CWT and STFT. Nonetheless, the consistency of CWT-derived findings across multiple independent models supports the robustness of our conclusions. Future studies with larger cohorts will be important for validating CWT-based OP metrics and establishing normative ranges. Specifically, we did not perform truly diagnostic analyses of the effects of using CWT vs STFT on PPSD after the energy preservation calibration (e.g., between-group effect sizes or discriminability of disease vs. control within each method), thus PPSD method comparisons are reported descriptively as CWT/STFT ratios of group means, without hypothesis tests or p-values. Within-recording dominant-frequency differences are reported as mean paired differences with 95% confidence intervals, without hypothesis tests. The synthetic validation employed idealized Gaussian-windowed bursts and autoregressive surrogates of recorded noise; real OPs are morphologically more complex. Finally, while this study focused on CWT and STFT to enable direct comparison of more commonly used methods, other adaptive time–frequency approaches, such as the HHT, warrant further investigation (23). However, the interpretability, computational efficiency, and strong physiologic grounding of CWT analyses make it particularly well suited for advancing OP analysis in both experimental and clinical electrophysiology.

## Conclusion

Our results demonstrate that CWT analysis using real Morlet wavelets provides a physiologically appropriate and analytically robust framework for characterizing OPs as compared to STFT. Under the evaluated analysis settings, CWT preserves the transient, nonstationary structure of OPs, produces more concentrated spectral representations, and improves temporal localization of OP features across experimental models and clinical settings. In synthetic signals of known content, CWT resolved closely spaced oscillatory events that the evaluated STFT implementation merged. Across cohorts, the descriptive CWT/STFT ratios of group mean PPSD ranged from 1.25 to 2.11, consistent with greater spectral concentration under equal-energy calibration. While both methods detected a significant dominant-frequency difference between RCS and BN rats, in the mouse and human cohorts, dominant-frequency differences reached significance with CWT but not STFT. Notably, these separate comparisons do not constitute a direct test of a difference in sensitivity between the methods. Nonetheless, taken together these findings support the adoption of CWT-based approaches for OP analysis as a complementary time–frequency method for assessing inner retinal function, with potential to improve the characterization and translational utility of ERG-derived biomarkers.

### Translational Relevance

Accurate and reliable quantification of OPs is critical for translating ERG recordings into reliable biomarkers of inner retinal function. This study demonstrates that CWT–based analysis provides a physiologically relevant time–frequency framework for characterizing transient OP activity that is variably captured by conventional Fourier-based approaches. By preserving time– frequency structure and reducing spectral spread within individual recordings, CWT improves the interpretability of OP time–frequency features across experimental and clinical settings, including models of retinal degeneration and patients with diabetic retinopathy. These properties are particularly relevant for monitoring disease progression, assessing therapeutic efficacy, and evaluating therapies affecting the inner retina, such as gene delivery, neuroprotective strategies, and optogenetics. Adoption of CWT-based OP analysis may therefore enhance the characterization and translational utility of ERG-derived functional measures, strengthening the link between preclinical studies and clinical assessment.

## Data and Code Availability

The complete analysis pipeline and the synthetic-validation data generators (MATLAB) are available at https://github.com/bgrosin/erg-wavelet-op (archived at https://doi.org/10.5281/zenodo.22983459) under an academic non-commercial license requiring citation; commercial use requires written permission (University of Pittsburgh Innovation Institute). No patient or animal recordings are distributed; the synthetic validation dataset is exactly reproducible from the included generators and noise-model parameter file.

