## Supplementary figures and images for "Wavelet Transform improves time-frequency characterization of the Electroretinographic Oscillatory Potentials as compared to Short-Time Fourier Transform across experimental and clinical models"

### Supplemental figure 1

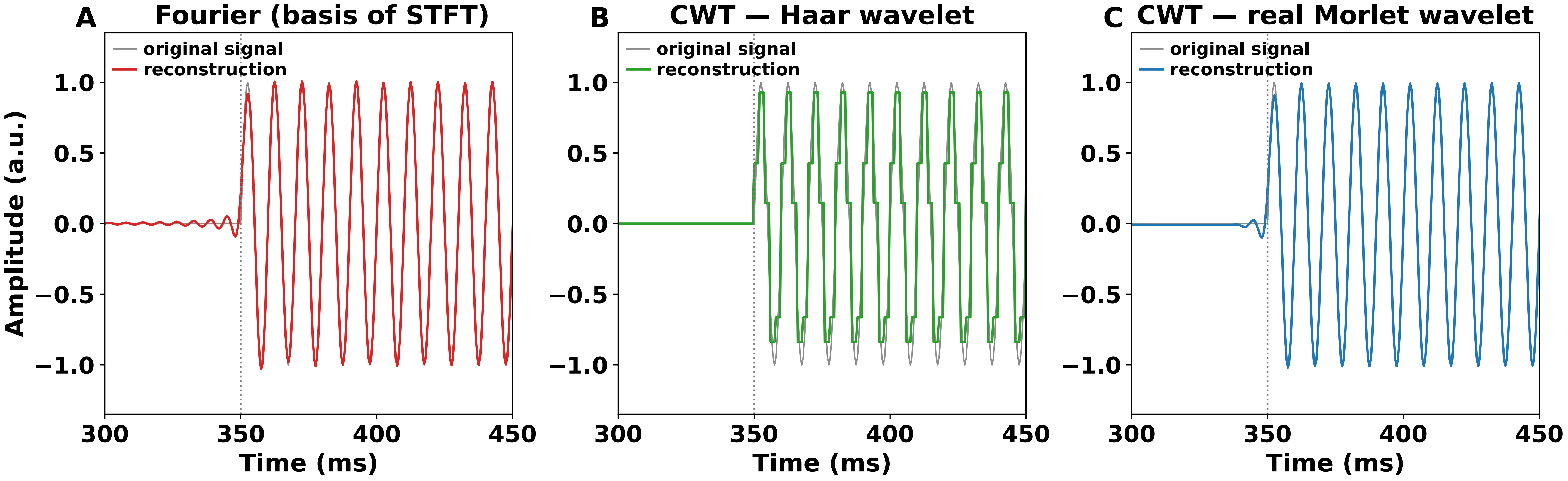
