## Supplemental figure 1 legend for "Wavelet Transform improves time-frequency characterization of the Electroretinographic Oscillatory Potentials as compared to Short-Time Fourier Transform across experimental and clinical models"

**Supplementary figure 1:**

Reduction of conspicuous Gibbs ringing by wavelet reconstruction. A 100 Hz sinusoid gated by a rectangular envelope with abrupt onset and offset (step discontinuities) was reconstructed from a band-limited (≤160 Hz) expansion using three bases and displayed around the onset transition (dotted line). (**A**) Fourier basis, which underlies the STFT: the reconstruction shows pronounced Gibbs ringing — a conspicuous train of oscillations flanking the discontinuity — characteristic of global, non-localized basis functions. (**B**) Continuous wavelet transform with the compactly supported Haar wavelet: the transition is reproduced essentially free of ringing, reflecting the strictly local support of the basis, although the waveform is approximated in a blocky, staircase fashion that poorly matches the oscillation. (**C**) Continuous wavelet transform with the real Morlet wavelet: the conspicuous ringing is markedly attenuated relative to the Fourier reconstruction while the oscillatory waveform is reproduced faithfully. Wavelet bases thus reduce the visible ringing associated with abrupt transitions, the degree of reduction depending on wavelet choice — greatest for compactly supported wavelets and more modest, though still substantial, for the real Morlet, which was selected for oscillatory-potential analysis because of its morphological resemblance to the signal (Fig. 1).
